# Stochastic Resonance Behavior of DNA Translocation with an Oscillatory Electric Field

**DOI:** 10.1101/2021.06.21.449299

**Authors:** Ining A. Jou, Rhys A. Duff, Murugappan Muthukumar

## Abstract

Stochastic resonance (SR) describes the synchronization between noise of a system and an applied oscillating field to achieve an optimized response signal. In this work, we use simulations to investigate the phenomenon of SR of a single stranded DNA driven through a nanopore when an oscillating electric field is added. The system is comprised of a MspA protein nanopore embedded in a membrane and different lengths of DNA is driven from one end of the pore to the other via a constant potential difference. We superimposed an oscillating electric field on top of the existing electric field. The source of noise is due to thermal fluctuations, since the system is immersed in solution at room temperature. Here, the signal optimization we seek is the increase in translocation time of DNA through the protein nanopore. Normally, translocation time scales linearly with DNA length and inversely with driving force in a drift dominated regime. We found a non-monotonic dependence of the mean translocation time with the frequency of the oscillating field. This non-monotonic behavior of the translocation time is observed for all lengths of DNA, but SR occurs only for longer DNA. Furthermore, we also see evidence of DNA extension being influenced by the oscillating field while moving through the nanopore.

## I. INTRODUCTION

The phenomenon of stochastic resonance is the synchronization between noise of the system and an external oscillatory field to optimize the response signal. Since the first studies on SR more than 30 years ago^1,2^, SR has been applied to many disciplines such as physics, chemistry, engineering, biology and biomedical sciences^3–10^. The early work of SR applies a weak periodic force to the noise-induced hopping of a particle in a bi-stable potential. The factors affecting particle dynamics are primarily the periodic force and the noise of the system while the energy barrier has a fixed height. SR has also been observed when other factors are incorporated in the system. For example, the use of uneven boundaries to create an entropic barrier instead of an energy barrier has been shown to lead to SR^11,12^. Investigation of SR of a flexible polymer has also been carried out using a combination of theory and simulations as the flexible chain crosses a bi-stable potential under an oscillatory force field^13,14^.

Another phenomenon related to noise of the system and particle motion that has garnered interest is resonance activation (RA). In resonance activation the cooperative interplay between the temporally modulated barrier and noise induced barrier crossing event can lead to a lowering of the crossing time. RA has been shown to be feasible for rigid particles^8,15^ and short polymers^16–19^. Nonetheless, the work we are presenting here is the result of SR and should not be confused with RA.

The SR of a flexible polymer chain has also been studied when the polymer undergoes translocation through a nanopore under an external periodic driving force. Here, the polymer is confronted with an entropic barrier, as it is subjected to a reduction in its conformational degrees of freedom due to the narrow pore^20^. A key result from the work of Mondal *et al*^21^ is that SR is not possible if the free energy barrier for polymer translocation has contributions only from chain entropy. However, when SR is achievable then the chain entropy can control the SR conditions significantly.

In biological systems SR has attracted special interest as thermal fluctuations play a major role in the dynamics of biomolecules. The application of SR offers a possible mechanism to take advantage of the stochastic movement of a biomolecule, and harness it to improve the signal output. One such particular interest is the application of SR in DNA translocation technology. The use of alternating electric field in a protein nanopore has been investigated using experiments and simulations^22–24^ but SR has yet to be observed on DNA translocating through a protein nanopore. The primary goal of this work is to investigate the feasibility of SR on a DNA translocating through a biological nanopore, and the dependence of the SR frequency on the length of DNA. The protein nanopore MspA has been chosen for this work for its well-reported structure and small sensing region^25–28^.

This work is organized as follows. The next section details the methodology with descriptions of the models and the simulation set-up. In Section III the results are presented and discussed. Section IV gives a brief overview and conclusions of this work.

## II. SIMULATION DETAILS

We use a coarse grained (CG) model of the protein nanopore MspA. The Protein Data Bank (PDB) entry used is 1UUN, with the additional residue mutations described by Butler *et al*^25^ performed on the CG model of MspA. These mutations remove the negatively charged residues at the constriction and allow for ease of DNA translocation through the nanopore. A schematic of the nanopore MspA is shown in Fig. 1. We use homopolymer DNA modeled using the united atom model^29,30^. The phosphates, sugars and bases are each represented with a bead of diameter 2.5 Å and mass of 96 Da. Only the bead representing the phosphate group has a charge *q* = −1*e*, where *e* is the elementary charge. We performed simulations for DNA of N = 23, 45, 90, and 120, where N is the number of bases.

**FIG. 1:**
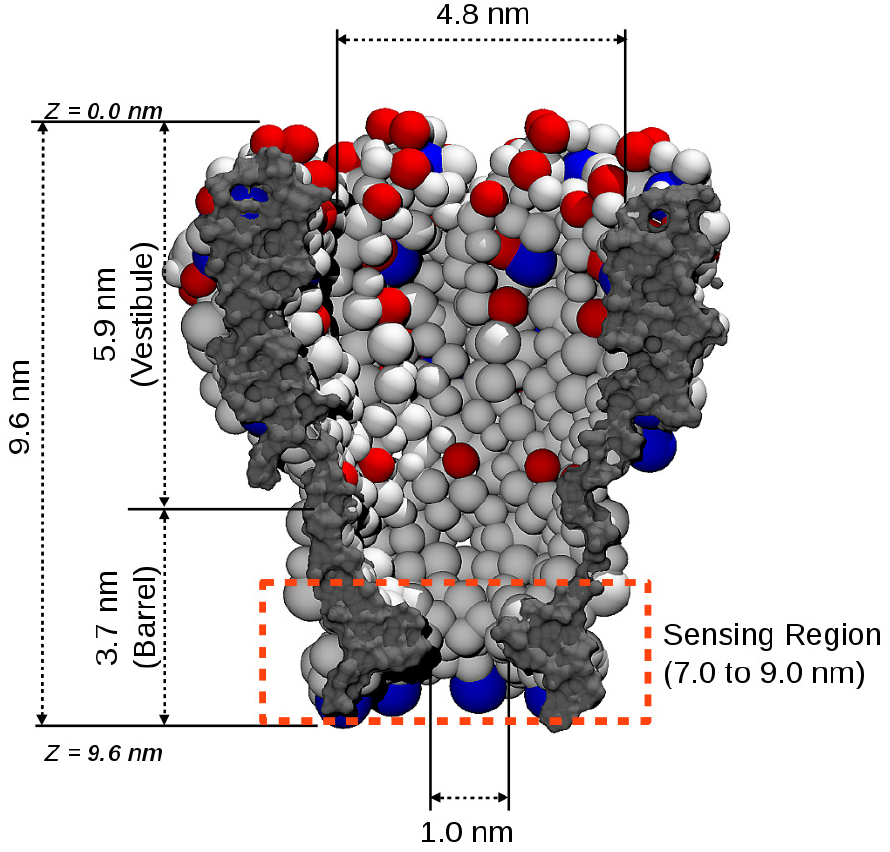
Schematics of the protein nanopore MspA. Blue beads each has a charge of +1*e*, red beads each has −1*e* charge and gray beads are neutral. The dashed rectangle indicates the sensing region where translocation time is calculated.

Langevin dynamics are performed to simulate the movement of DNA through the system. Through this approach the collision with water molecules is replaced with a stochastic force which represents the total net force due to the water molecules at a given time. The nanopore and the membrane are treated as a rigid body fixed in space so interactions among beads of the nanopore and membrane are not considered. Simulations of DNA undergoing translocation through the protein nanopore in the presence of an external electric field are performed using LAMMPS^31^ package. The position of the DNA is updated using Langevin dynamics of the *i*-th bead of the DNA:

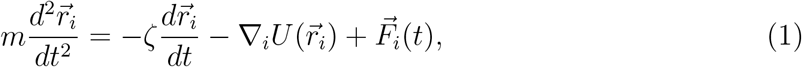

where *m* is the mass of each DNA bead, 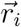 is the position of the *i*-th bead, *ζ* is the friction coefficient, *U* is the total potential energy acting on the *i*-th bead, and 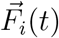 is the stochastic force acting on the bead. The random force term 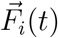 is the net force due to the solvent molecules acting on the *i*-th bead at time *t*, satisfying the fluctuation-dissipation theorem 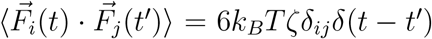, where *k_B_* is the Boltzmann constant, *T* = 298 K is the temperature, and *δ*(*t*) is the Dirac delta function. The value of the friction coefficient sets the time scale for the dynamics. In this work the friction coefficient was fixed arbitrarily as 1 in simulation reduced unit for simplicity since we do not know the drag coefficient on each nucleotide in reality. As a result, the time scale is arbitrary in the Langevin Dynamics simulations. The time scale in real units can be obtained by comparing with experiments^30^.

The total potential energy *U* acting on the *i*-th bead can be written as follows:

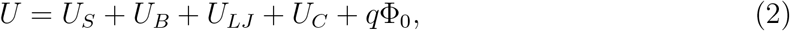

where *U_S_*, *U_B_*, *U_LJ_*, *U_C_* and *q*Φ_0_ are the different interaction potentials present in the system, as described below.

The spring bonding potential, *U_S_* = *k_s_*(*r_s_ − r*_0_)^2^, acts between connected beads, where *k_s_* = 171 kcal/mol*·*Å^2^ is the spring constant^32^, *r_s_* is the distance between the *i*-th bead and its adjacent bead, and *r*_0_ = 2.5 Å is the equilibrium distance. The bending angle potential, *U_B_* = *k_a_*[*cos*(*θ*) *− cos*(*θ*_0_)]^2^, is between the side chain and the backbone, where *k_a_* = 60 kcal/mol is the bond angle constant^33^, *θ* is the angle between the side chain and the phosphate backbone, with *θ*_0_ = 65° being the equilibrium bond angle^33^. The Lennard Jones (LJ) potential modeling the excluded volume interactions is 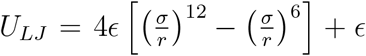, where *ϵ* = 0.5 kcal/mol, *σ* = 2.5 Å, and *r* is the distance between the *i*-th bead and the *j*-th bead of the system. The LJ potential is truncated at *r* = *r_c_*, where *r_c_* = 1.12*σ*. The screened electric interaction, 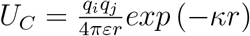, is modeled using the Debye-Hückel potential, where *q_i_* and *q_j_* are the charges of the *i*-th and *j*-th beads in the system, *κ* = *L_D_^−^*^1^ = 0.182 Å^−1^ is the inverse Debye length in 0.3 M KCl solution, *ε* = *ε*_0_*ε_r_* is the permittivity, *ε_r_* = 2 in the protein and the lipid bilayer and *ε_r_* = 80 everywhere else, and *ε*_0_ is the permittivity of free space. The simulation starts with DNA partially inside the vestibule cavity to reduce unnecessary simulation time during the capture stage, since the main focus of this work is to understand the translocation dynamics of DNA through the nanopore and not capture of DNA.

An external potential difference, *V_e_*, of 180 mV is applied between the *cis* and *trans* chambers of the system. The electrostatic potential of the system as a result of the applied potential, charges in the nanopore, and ions in the solution is calculated using the Poisson-Nernst-Plank [PNP] formalism^34–37^. The last term of Eq. 2 is the electrostatic potential of the system, where *q* is the charge of the *i*-th bead and Φ is the electrostatic potential through the pore computed by using the PNP procedure, which couples the steady state Poisson equation with the Nernst-Planck equations. This is done by solving self-consistently the Poisson equation:

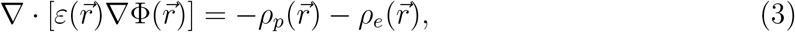

together with the steady state Nernst-Planck equation:

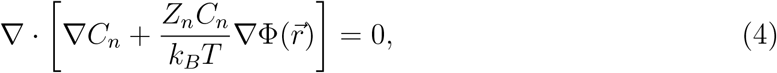

to obtain the local concentrations of the cations and anions following similar procedures as discussed in literature^29,38^. In Eq. 3 the *ε* is as described before and its value differ depends if in solution or in biomolecule. In Eq. 4, *n* = *{K*^+^*, Cl^−^}* are the ions in the solution, and *Z_n_* = 1*e* or −1*e* depending on the ion species. The charge densities in Eq. 3 are due to all the fixed charges in the system such as the charges on the protein, *ρ_p_*, and the mobile ions in the solution, *ρ_e_*, where 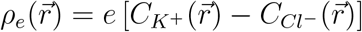. The electrostatic potentials are calculated using Eqs. 3 and 4 along the center of the nanopore at (0, 0*, Z*).

DNA starts partially inside the entrance of the nanopore at *z* = 2.0 nm and allow to relax in the absence of the external applied potential difference *V_e_*. The relaxation time of DNA is set to be long enough to allow its gyration radius to reach an equilibrated length. After relaxation, the external potential difference *V_e_* and the oscillating electric field are applied and the trajectory of the DNA through the nanopore is simulated. There are at least 1500 simulations performed for each frequency.

In addition to the existing electric field from Φ_0_, we superimposed an oscillating electric field inside the nanopore of the form

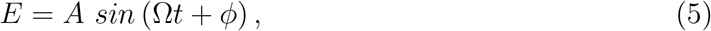

where *A* = 0.008323, Ω is frequency, *t* is time, and 0° ≤ *ϕ* ≤ 360° is a randomly generated phase angle. The amplitude *A* is equivalent to considering a potential difference of 50 mV in a nanopore of length 95 Å. The various values of Ω in this work are expressed as Ω = *nω*, where *ω* = 0.757 *×* 10^−3^ *rev/au*, *n* is a multiplying factor and *au* is the arbitrary simulation time unit. Fig. 2 shows examples of the superimposed voltages. The randomly generated phase angle is included to mimic the condition in experiment where DNA does not enter the nanopore at the same point during the oscillation. This field is added to the electric field from the PNP calculation as a simple approximation of the electric field inside the nanopore region.

**FIG. 2:**
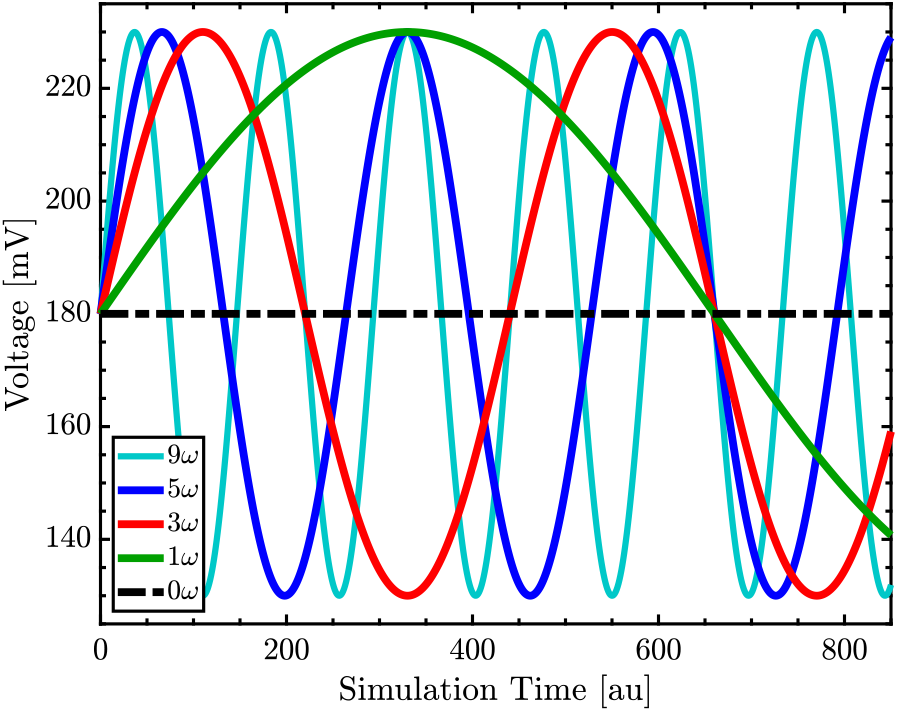
Voltages of different frequencies during simulation time. The black dashed line in the middle is the zero frequency case.

The frequencies in simulation units can be mapped onto real units using the following relationship:

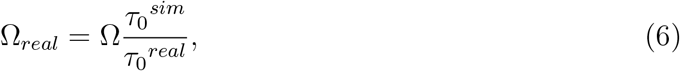

where Ω_*real*_ is the frequency in real units, *τ*_0_^*sim*^ is the translocation time in *au* when no oscillating field is applied, and *τ*_0_^*real*^ is the translocation time in real units when no oscillating field is applied. Specific example will be discussed in the next section.

## III. RESULTS & DISCUSSION

Let us consider first the electrostatic potential when no oscillating field is applied to the system. The geometry and charge distribution of the MspA nanopore produces a unique electrostatic potential profile through the pore as shown in Fig. 3. The electric potential in MspA nanopore rises very slowly within the vestibule region, up to about 40 mV potential difference at just before the sensing region. For a negative charge moving in the vestibule, the maximum electric energy difference in this region is about 1 kT at room temperature, the same as thermal fluctuations. DNA dynamics will be mostly diffusive in the vestibule region due to the low electric energy difference. The electric potential then quickly rises within the sensing region until the maximum potential difference of 180 mV is reached just outside the pore. This potential difference of about 140 mV within 20 Å inside the sensing region produces a very strong electric field (7 mV/Å) to drive DNA to translocate.

**FIG. 3:**
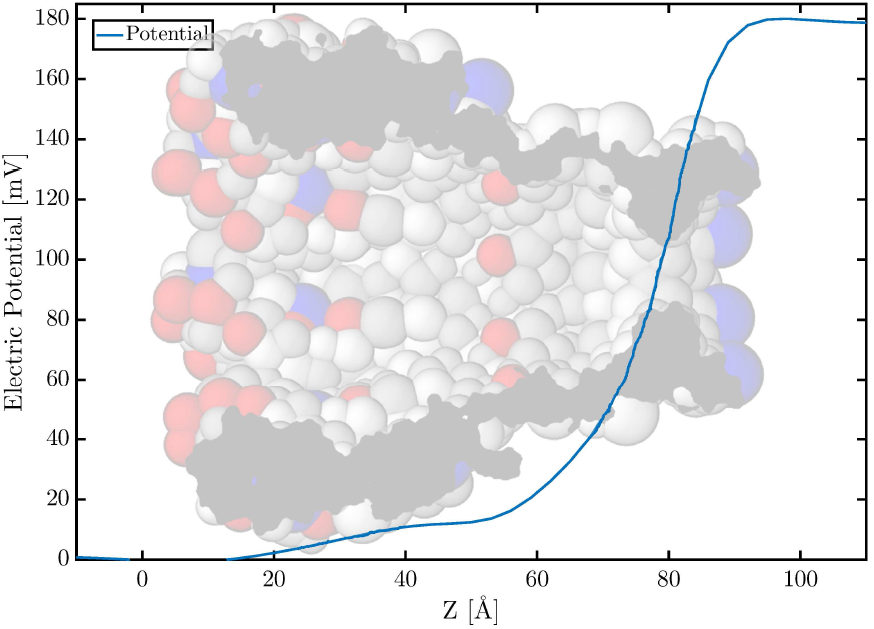
Electrostatic potential through the nanopore along the *z*-axis calculated using the PNP formalism.

When the oscillating field is applied, it is superimposed over the existing electric field inside the nanopore. The oscillating field can increase the existing electric field to strongly facilitate DNA motion, or it can decrease the existing field and hinders DNA translocation. The oscillation frequency determines how often the existing electric field is increased or decreased. We say the system is in a *facilitating* state when the phase of the oscillating field increases the existing electric field since DNA translocation becomes strongly facilitated. When the oscillating field is in a phase that decreases the existing electric field then we say the system is in a *hindering* state since DNA translocation is slowed down. The duration of each of the states is one half the period of the oscillating field. The specific values of the half period of each frequency are shown in Table I.

**TABLE I:** The half period of each frequency. The half period corresponds to the duration of the *facilitating* or *hindering* state.

| Frequency [$\omega$] | Half Period [au] |
| --- | --- |
| 0.1 | 6600 |
| 0.5 | 1321 |
| 1 | 661 |
| 2 | 330 |
| 3 | 220 |
| 4 | 165 |
| 5 | 132 |
| 6 | 110 |
| 7 | 95 |
| 8 | 83 |
| 9 | 74 |
| 10 | 66 |
| 20 | 33 |

The translocation process starts when DNA enters the sensing region without reverting to the vestibule and completes a successful translocation. Translocation time, *τ*, is defined as the time difference between when at least one nucleotide has entered the sensing region, and when all nucleotides have exited the sensing region at *z* > 9.0 nm. At any point during translocation if all the DNA bases retreat completely out of the sensing region to the vestibule then the translocation timer is reset to 0.

### A. Translocation Dynamics of DNA (N=90)

During the translocation stage DNA enters the sensing region then exits the barrel on the *trans* side. The probability distribution of the translocation time P(*τ*) in the presence of various oscillating frequencies is shown in Fig. 4. At zero frequency, the near-constant electric field in the sensing region provides the primary driving force for DNA translocation since the electric field in the vestibule region produces minimal drift on the DNA. The resulting normal distribution of *τ* has a clear defined peak position at 807 *au* with a relatively small spread as shown in Fig. 4 for 0*ω*. This distribution resembles the probability distribution of first passage time with one absorbing boundary and drift where the boundary is at a distance away from the starting point^39^.

**FIG. 4:**
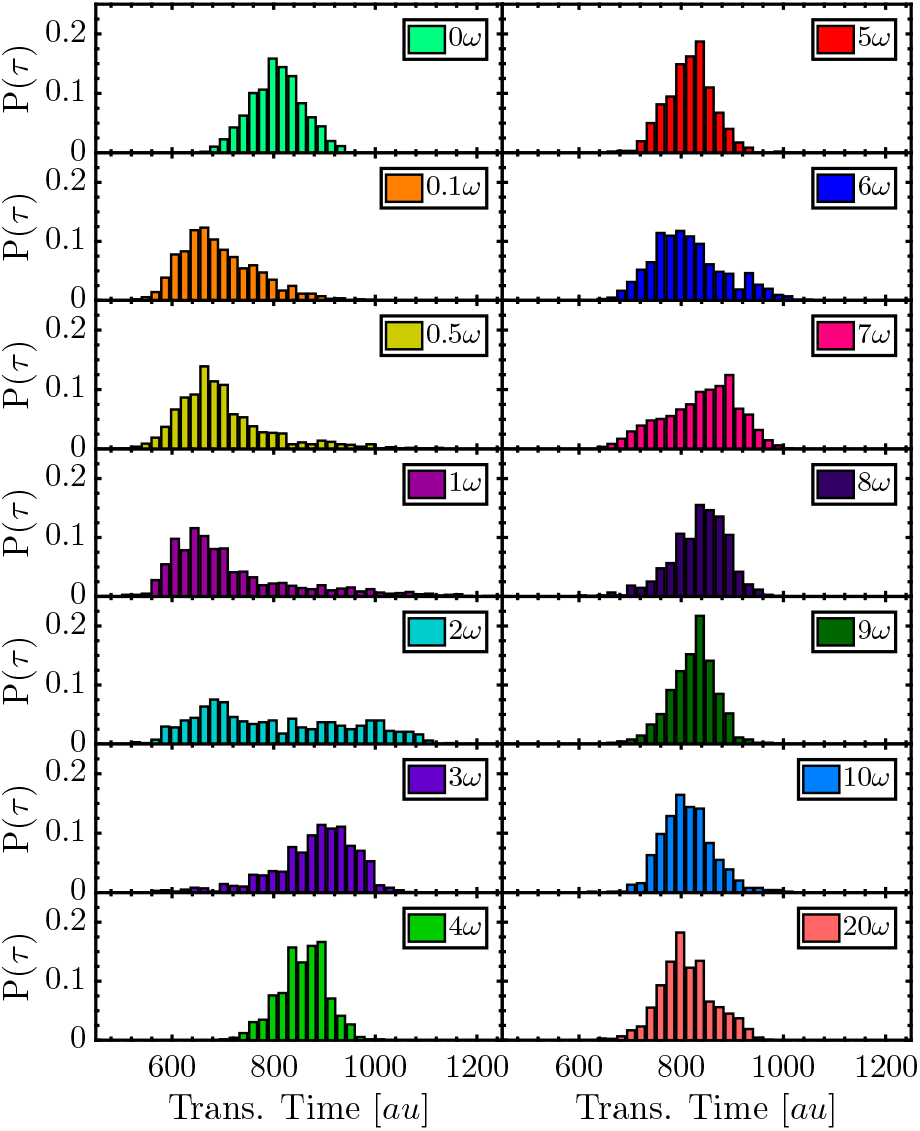
Probability distribution of DNA translocation time in the presence of oscillating field at various frequencies when N=90. Translocation time is defined as the time taken by DNA to move through the sensing region.

For frequencies 0.1, 0.5 and 1*ω* the probability distribution is shifted towards the left, with a peak around 690 *au*. This means that the successful translocations took less time than the zero frequency case. At these frequencies the period of oscillation is much longer than the typical translocation time (see Table I), resulting in duration of the facilitating or hindering state also being longer than the translocation time. If DNA enters the pore while the oscillating field is in the facilitating state then the translocation process is facilitated. When translocation is facilitated the time it takes for DNA to go through the nanopore is shorter. We confirmed this by plotting the translocation time of 0.1*ω* with the initial phase angle *ϕ* as shown in Fig. 5. As mentioned before, *ϕ* is the phase at which the DNA is first inside the nanopore. The value of *ϕ* is randomly generated between 0° and 360° to mimic in experiment where DNA could enter the nanopore at any phase angle. Analysis of *ϕ* along with the duration of facilitating/hindering state allows us to understand how many of facilitating or hindering state did the DNA experience during translocation. In Fig. 5 the phase angles between 0° to 180° would put the system in a facilitating state, since the oscillating field is described by a sine function. The translocation times directly reflect the amplitude of the initial phase when frequencies are very low.

**FIG. 5:**
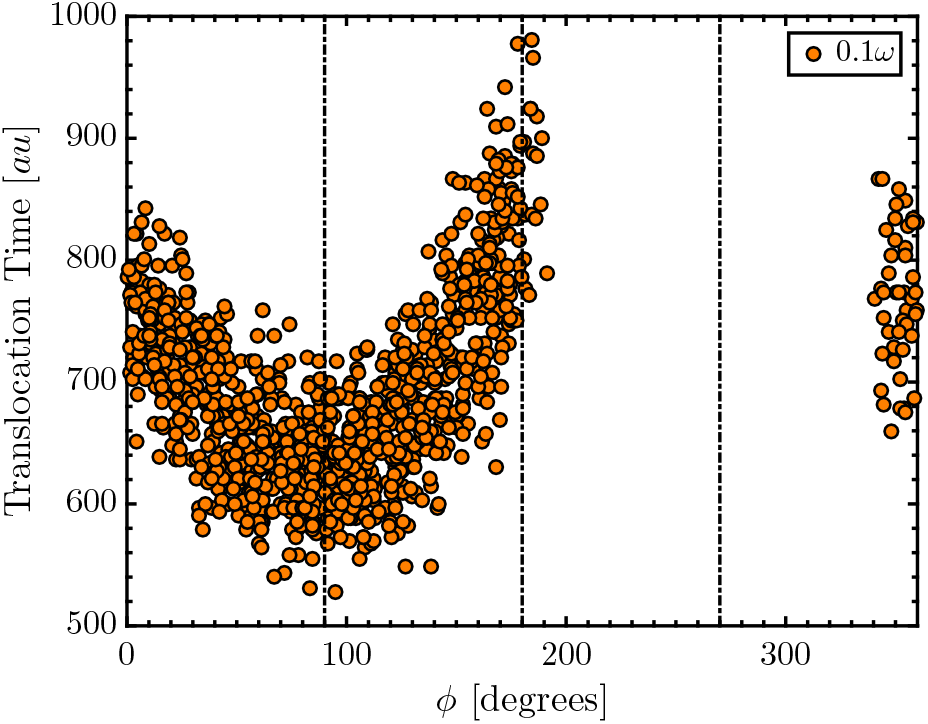
Scatter plot of DNA translocation time *vs.* the phase *ϕ* at *t* = 0, *i.e.* the phase that DNA encounters when it enters the nanopore. Vertical dashed lines are at 90°, 180°, and 270°. This is for N=90 at frequency 0.1*ω*.

However, not all values of *ϕ* result in translocation. In Fig. 5 there is very few successful translocations for values of *ϕ* between 180° and 360°. If the system is in the hindering state when DNA enters the pore, translocation is not likely to occur as there is not enough driving force to push DNA into the sensing region. For frequencies 0.5 and 1*ω* translocation times are becoming longer if DNA started translocation towards the end of the facilitating state (see Fig. S3(a) in Supplementary Information).

At frequency equals to 2*ω* the duration of facilitating or hindering state is about 330 *au*. This means that during the translocation stage DNA will experience at least once of each the hindering state and the facilitating state. DNA translocation dynamics is facilitated then hindered for approximately the same amount of time, or *vice versa*. Consequently, the translocation time distribution shows a wide spread with no clearly defined peak. One can argue a small peak is visible at around 660 *au*. Looking at the phase angles in Fig. S3(a) for 2*ω*, it is easy to see that these faster translocations correspond to DNA that started its translocation process when the facilitating state just started.

For frequency 3*ω* the translocation time distribution has shifted towards the right past the peak of zero frequency case, with a peak at around 902 *au*. At this frequency, the duration of facilitating/hindering state of 220 *au* is approximately 1/4 of the peak translocation time. A translocating DNA would experience the facilitating and hindering states about 2 times each. As shown in Fig. S3(a) for 3*ω*, most of the translocation occurs if DNA enters the nanopore when the facilitating state has just started or when the hindering state in ending. This is also the first frequency where the length of the translocation time no longer shows a clear increase or decrease with the initial phase. On the other hand, how often DNA experiences the facilitating/hindering state has a dominant effect on the translocation time. What this mean is that the translocation time is not so much affected by the initial phase, but more so by the frequency of the oscillating field. Nevertheless, initial phase can still influence the translocation being successful or not.

For frequencies 4*ω* to 10*ω* the peak location is shifted slightly towards the left again to around 810 *au*. There is no significant change to the distribution besides the slight sifting of the peak position around the zero frequency case. As the duration of facilitating/hindering state becomes shorter with increased frequency, DNA has less time to react to the oscillating field. In Fig. S3(a), we can see that translocation times are recorded for wider and wider range of *ϕ*. This follows as the duration of facilitating/hindering state decreases with frequency, *ϕ* become less important. At the highest frequency 20*ω* DNA displays similar distribution to the zero frequency case. Translocation is recorded for all values of *ϕ* and no longer plays a significant role in DNA translocation.

The shifting back and forth of the peak of *P* (*τ*) becomes less evident for frequencies 4*ω* to 10*ω*, as shown in Fig. 4. At these frequencies, the shape of the *P* (*τ*) changes slightly in different frequency while keeping the peak position at around 800 *au*. As the duration of facilitating/hindering state becomes shorter with increased frequency, DNA has less time to respond to the changing force. DNA eventually behaves similarly to the zero frequency case when the oscillation frequency is too high.

#### 1. Stochastic Resonance Behavior

From the probability distribution of translocation time in Fig. 4 we saw that the translocation time is affected by the applied frequency through the duration of the facilitating/hindering state. Specifically, we have plotted the mean translocation time against frequency in Fig. 6 to show the non-monotonic behavior of mean translocation time with frequency. The mean translocation time of the zero frequency case is 807 *au* which coincides with the peak location of its distribution in Fig. 4. For frequency 10 and 20*ω* the mean translocation time is almost the same as the zero frequency case at 808 and 806 *au*, respectively. This is as expected since at higher frequency the duration of each facilitating or hindering state is very short for DNA to react to.

**FIG. 6:**
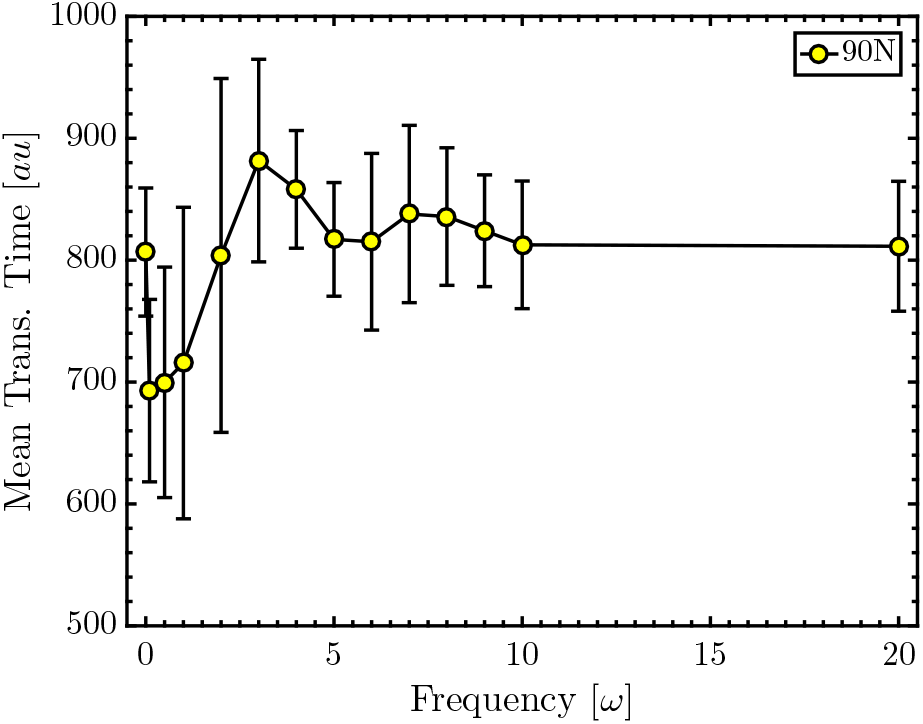
Mean translocation time of DNA with N = 90.

The drop in the translocation time from 0.1 to 1*ω* is due to the prolonged effect of the facilitating state as discussed before, where the entire translocation process is during the facilitating state. At 2*ω* the mean translocation time is comparable to the zero frequency case, although its time distribution in Fig. 4 is very different from the zero frequency case. At 3*ω* the mean translocation time is maximum at about 10% longer than the zero frequency case. This shows that at this specific frequency the effect from the oscillating field on DNA dynamics is tuned in to the motion of DNA in such a way to notably increase the translocation time. The SR frequency for N=90 is then at 3*ω*.

As frequency increases beyond 3*ω* the mean translocation time decreases then increases slightly with frequency. The difference between the zero frequency case and these translocation times are 4% or less. At very high frequency, the translocation time for zero frequency case is recovered.

### B. Dependence of SR on DNA Length N = 23, 45, 90, and 120

We performed simulations on three other lengths of DNA and the mean translocation times per base are plotted in Fig. 7. The probability distribution of translocation time of these DNA lengths are included in Fig. S1 in the Supplementary Information, along with the scatter plots of the translocation time with the initial phase in Fig. S2 and S3. The translocation time is normalized by the number of bases to compare the different DNA lengths.

**FIG. 7:**
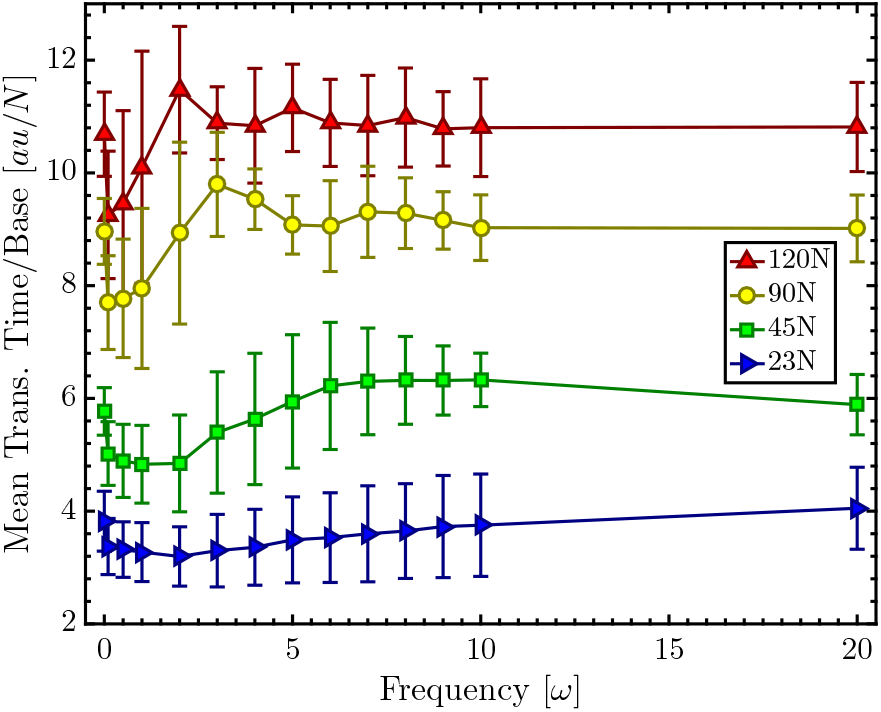
Mean translocation per base *v.s.* frequency for different DNA lengths.

A non-monotonic behavior in the mean translocation time is seen for all DNA length. At some very low frequency, translocation times of all 4 DNA display a minimum. This drop in time is an effect of the prolonged facilitating state and is more prominent for longer DNA. After the minimum the translocation time gradually increases with frequency.

For longer chains, N=90 and 120, a maximum is clearly seen in the mean translocation time at 3*ω* for N=90 and 2*ω* for N=120. We say that SR is achieved for these DNA at those specific frequencies. We do not see a clear maximum for either the N=23 and N=45. This shows that the SR frequency is directly related to the length of DNA.

The relation between translocation time and DNA length for different frequencies is shown in Fig. 8. For the lengths of DNA studied, a linear dependence is shown between the mean translocation time and the DNA length for most frequencies. As expected, the results of higher frequencies such as 20*ω* are very similar to that of the zero frequency case. At lower frequency, such as 0.1*ω*, the translocation time is lower than the zero frequency case but still shows the same linear increase with DNA length. An exception to the linear behavior is at 2*ω* and 3*ω*, where a cross over from lower to higher translocation time occurs. At 3*ω* the cross over is around N=60, which could be an indication of the shortest DNA length to observe stochastic resonance at this frequency. At 2*ω* the cross over is not until after N=90.

**FIG. 8:**
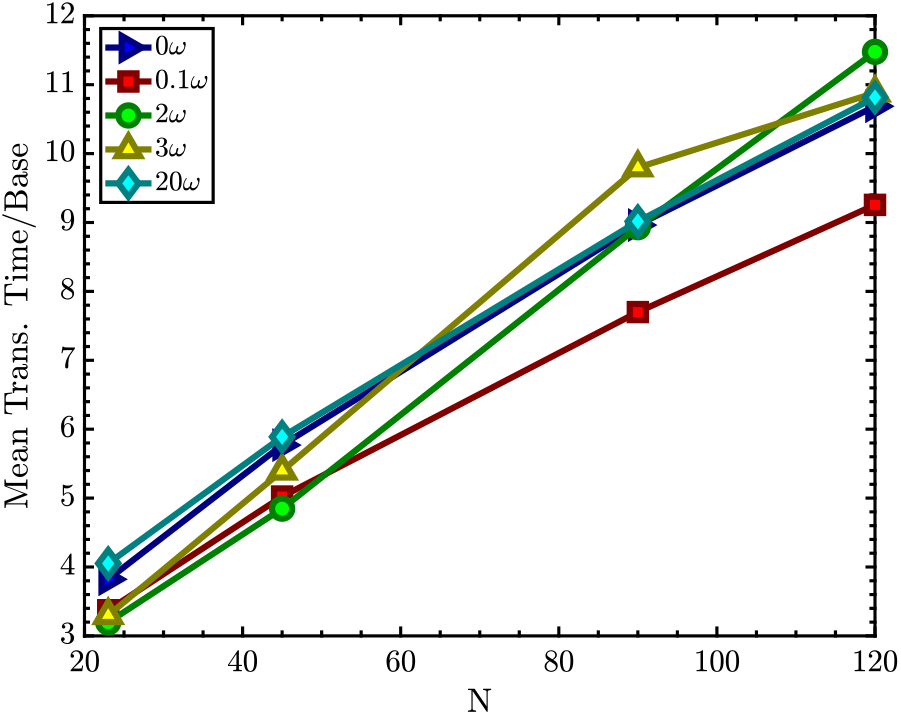
Mean translocation per base *v.s.* DNA length at different frequencies.

To compare our results to experiments, the mean translocation time in zero frequency, *τ*_0_, can be plugged into Eq. 6 to estimate the frequency in real units from simulation units. For example, we use the case of 90N where Ω = 3*ω* and *τ*_0_ = 882 *au* to estimate the resonance frequency in real units. Assuming that the mean translocation time in zero frequency in experimental condition is 5 ms, from Eq. 6 we get Ω_*real*_ = 355 Hz.

### C. Local Dynamics along the DNA

In order to see exactly how the oscillating field is affecting the local dynamics of DNA, we looked at the trajectory of each DNA base as they move through the nanopore. This type of data is not accessible in experiments and is currently only available through simulations. The *z*-coordinate of each DNA base is plotted against the simulation time. As time advances each DNA base moves from the entrance of the nanopore at *z* = 0 Å towards the exit of the nanopore at *z* = 90 Å. From frequencies 0, 3, and 10*ω* one sample trajectory of each frequency is shown in Fig. 9. The corresponding oscillating voltage of each frequency is shown on the top for each plot. Since N=90 there are 90 trajectories in each figure, where the darkest blue is the first DNA base and the darkest red is the last. The solid black line in the middle of the color lines is the center of mass trajectory. The vertical dashed line indicates when DNA enters the sensing region and marks the start of the translocation time. The time transpired before the vertical dashed line is the waiting time. The two horizontal dotted lines show the *z*-coordinates of the sensing region.

**FIG. 9:**
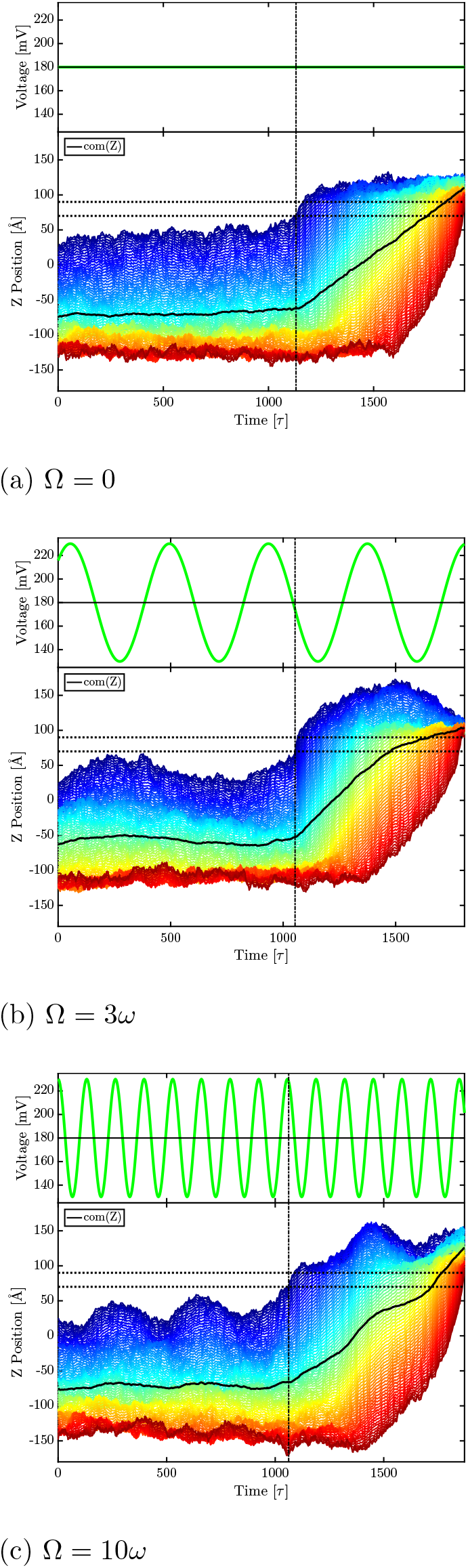
The *z*-coordinate of each DNA base *vs.* simulation time in one complete translocation simulation. There are 3 different frequencies shown here, with the corresponding oscillating voltage for each frequency. The DNA has N=90 so there are 90 lines from the darkest blue to the darkest red, where each line shows the *z*-trace of each base. The darkest blue is the *z*-trace of the first DNA base and darkest red is the *z*-trace of the last base. The space between the two horizontal dotted lines corresponds to the sensing region. The vertical dash line indicates the beginning of the translocation process

First, let us consider the zero frequency case as shown in Fig. 9(a). There is oscillating voltage here so in the voltage plot at the top of Fig. **??**(a) is just a flat line representing the constant applied potential difference. Inside the vestibule DNA extends slowly as the separation between bases increases slowly, visible from the separated blue lines. For the portion of the DNA that are still outside the nanopore (orange and red lines) individual base trajectories are not distinguishable, because outside the nanopore DNA can move in the *x* and *y* directions as well. The constant extension inside the pore is due to the geometric confinement, and the slow gradual extension is due to the low electric field in the vestibule. The DNA extension increases rapidly as DNA goes through the sensing region, since this is the narrowest part of the nanopore with a strong electric field. As soon as DNA exits the nanopore they are free to move in the *x* and *y* directions so the lines become indistinguishable.

When the oscillating frequency is 3*ω* the extension of DNA is no longer steady as shown in Fig. 9(b). The top of Fig. 9(b) shows the oscillating voltage (green line) on top of the constant voltage at V=180 mV. For this particular translocation the initial phase was around 70*◦* so DNA first encountered facilitating state which helped it to move further into the nanopore. As the facilitating state reaches maximum the DNA extension inside the nanopore also reaches maximum at a short time later. As the oscillating voltage enters the hindering state the DNA extension is also reduced after a short delay. Upon entering the sensing region DNA is rapidly extended and exited the nanopore. Comparing to the zero frequency case, we can see a further DNA extension even after exiting the nanopore as a consequence of the last facilitating state inside the nanopore.

At 10*ω* as shown in Fig. 9(c) the facilitating and hindering state have very short duration of 66 *au*. DNA shows a delayed extension/contraction loosely corresponding to the oscillation while the movement of its center of mass is still comparable to that of other frequencies in Fig. 9(a) and (b). After exiting the nanopore further extension/contraction can also be seen corresponding very loosely to the oscillating field.

Aside from the stochastic resonance in translocation time, there is an additional synchronization effect between the oscillating field and the DNA extension as the number of peaks in Fig. 9 scales loosely and qualitatively with frequency. A perfect synchronization between DNA extension and the frequency will not be possible as DNA moves sluggishly due to its chain length and thermal fluctuations.

### D. Waiting Stage Dynamics (N=90)

From simulation data we are able to separate the entire translocation process into two stages: the waiting stage and the translocation stage. The waiting stage is when DNA is inside the vestibule before entering the sensing region. Waiting time, *τ_W_*, is then the time DNA spent in this stage. During the waiting stage part of DNA is in the vestibule region and part of it still outside the nanopore. During this time, DNA movement inside the vestibule is mostly random motion due to the low electric field inside the vestibule. As such, the waiting time distribution without the oscillating force has a wide spread in Fig. 10 for 0*ω*.

**FIG. 10:**
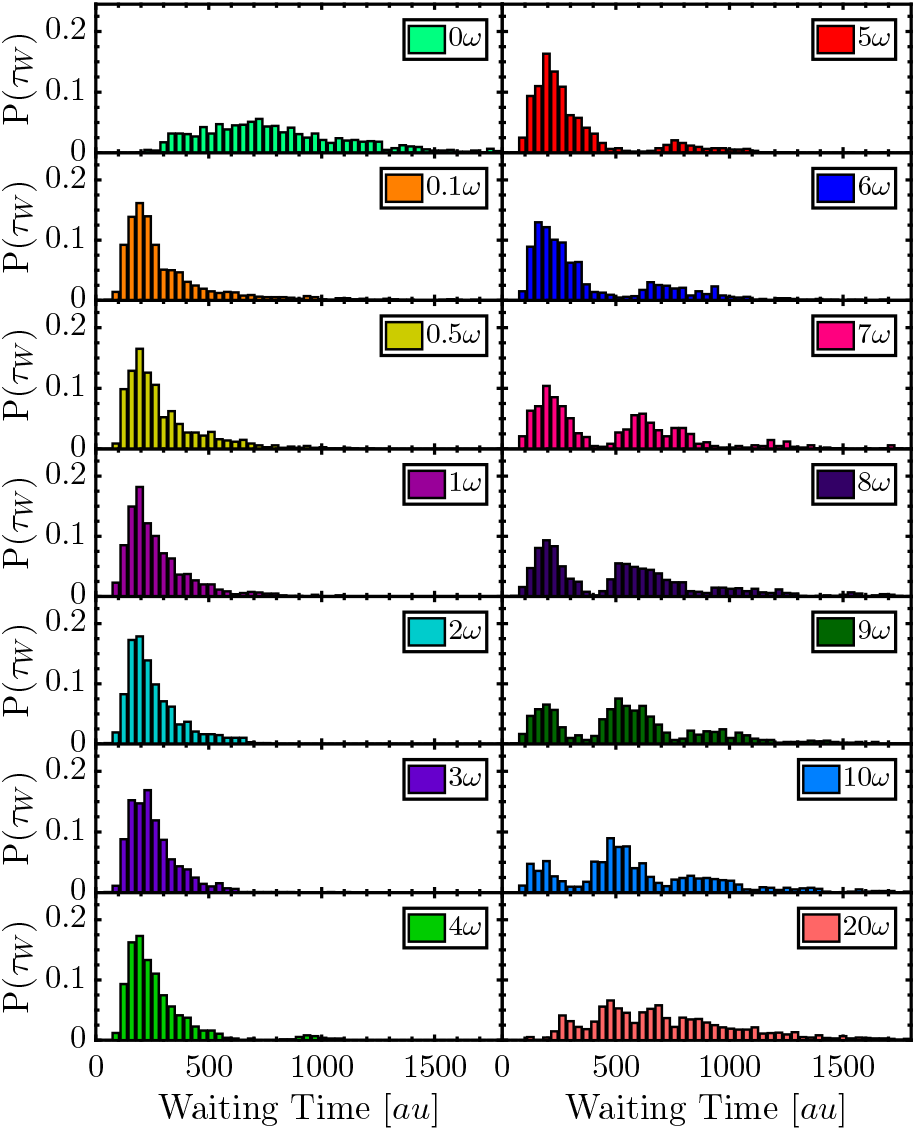
Probability distribution of DNA waiting time in the presence of various frequencies of the oscillating field when N=90. Waiting time is the time DNA spent above the sensing region before translocation starts.

For frequencies 0.1*ω* to 4*ω* the waiting time distribution is shifted towards the left, with a peak position around 200 *au* and small time difference among the different frequencies. Considering that these are the waiting times of only successful translocations, we know that these translocations occur mostly when the DNA enters the nanopore while the system is in the facilitating state. As the duration of the facilitating state at these frequencies range from 165 to 6600 *au*, DNA is facilitated long enough to pass to the sensing region.

At frequencies 5*ω* to 20*ω*, in addition to the first peak, a second and even a third peak appear in the distribution. These peaks show that within the same frequency, there is more than one most likely time for DNA translocation. The first peak is around 190 *au*, similar to the previous lower frequencies. The waiting time in the first peak is when the DNA experienced mostly facilitated state while it is in the vestibule. As frequency is increased, this first peak decreases and gives rise to a secondary peak. In frequencies 7*ω* to 10*ω* a third peak also appears. Considering that thermal fluctuations is comparable to the strength of the oscillating field, this means that DNA is also likely to randomly move in the vestibule, not necessary towards the sensing region. It is possible for DNA to not arrive in the sensing region during the first facilitated state, and manages to stay the vestibule during the hindering state. If DNA stays in the vestibule through this time then DNA will be facilitated towards the sensing region when the next facilitated state cycle through. The strip patterns in the waiting time *v.s.* phases in Fig. S6(a). The strips in the plot reflects the peaks seen in Fig. 10. The additional insight we get from Fig. S6(a) is that DNA can start from similar facilitating state but not arrive the sensing region in the same time.

At very high frequencies, such as 20*ω*, the duration of each facilitated and hindered state is so short that DNA does not have time to react to it. The dynamics of DNA start to resemble the zero frequency case as the peaks in the distribution of waiting times become very close together in Fig. 10 and the separation between stripes in Fig. S6(a) becomes insignificant.

### E. Waiting Time of Different DNA Length N = 23, 45, 90, and 120

Overall, DNA mean waiting time varies non-monotonically with frequency as shown in Fig. 11. The zero frequency case shows the largest mean waiting time for all lengths of DNA. In the presence of the oscillating field the waiting time is lowest at 0.1*ω* for shorter DNA, N=23 and 45. As for longer DNA, N=90 and 120, the lowest waiting time is at about 2*ω*. With the oscillating field applied, DNA motion is facilitated and the mean waiting time decrease by more than 50% at lower frequencies. As frequency increases, DNA exhibits a non-monotonic increase in waiting time. Specifically, for longer DNA the waiting time from 0.1*ω* to 4*ω* shows no significant variation. These are the waiting times with one peak in the probability distribution. From 5*ω* and on wards the slope of the waiting time is bigger, as some waiting times are becoming longer even within the same frequency. These are the frequencies with the appearance of secondary and tertiary peaks in the waiting time distribution. The waiting time distribution of other length of DNA is in Fig. S4 and the scatter plots of the waiting time with the initial phase in Fig. S5 and S6.

**FIG. 11:**
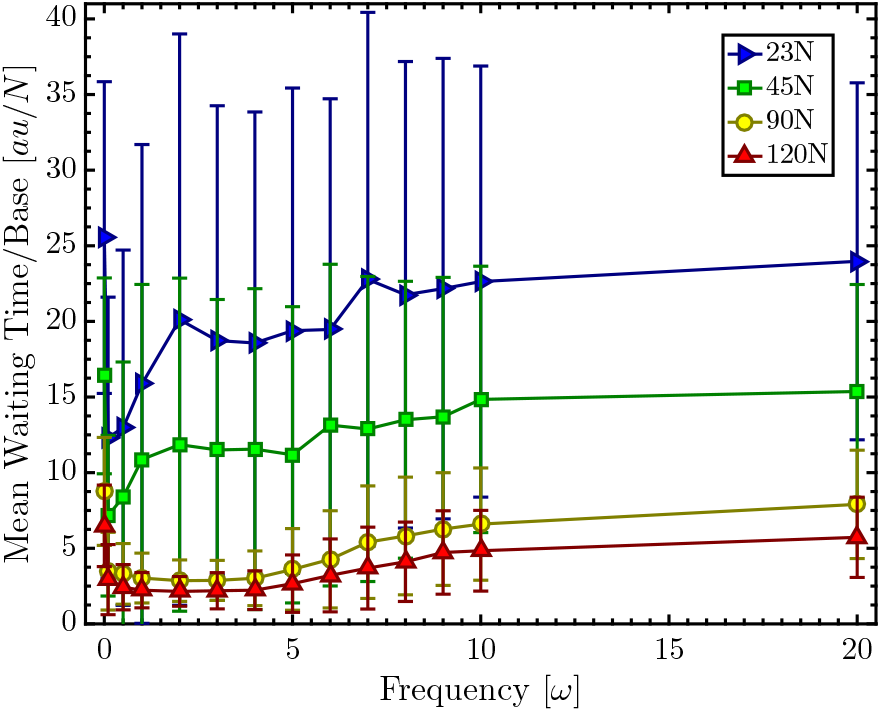
Mean waiting time per base *v.s.* frequency for different DNA lengths.

Note that for the same frequency, the mean waiting time decreases as DNA length become longer as opposed to the translocation time being proportional to the chain length. We show in Fig. 12 that the waiting time decreases with the chain length. We can see that for very long DNA the waiting will all be short and similar. This counter-intuitive decrease in waiting time with chain length originates from the local entropic push that polymer in a region with a much higher number of monomers exerts on the region with very low or no monomers^40^. For long DNA inside the vestibule, there is a large number of monomers compared to no monomers inside the sensing region, so at the entrance of the sensing region some monomers are entropically pushed into the sensing region. Due to the high electric field inside the sensing region, the electric force on just a few monomers is sufficient to initiate the translocation process. For shorter DNA, the number of monomer inside the vestibule is not high enough to force its monomers into the sensing region.

**FIG. 12:**
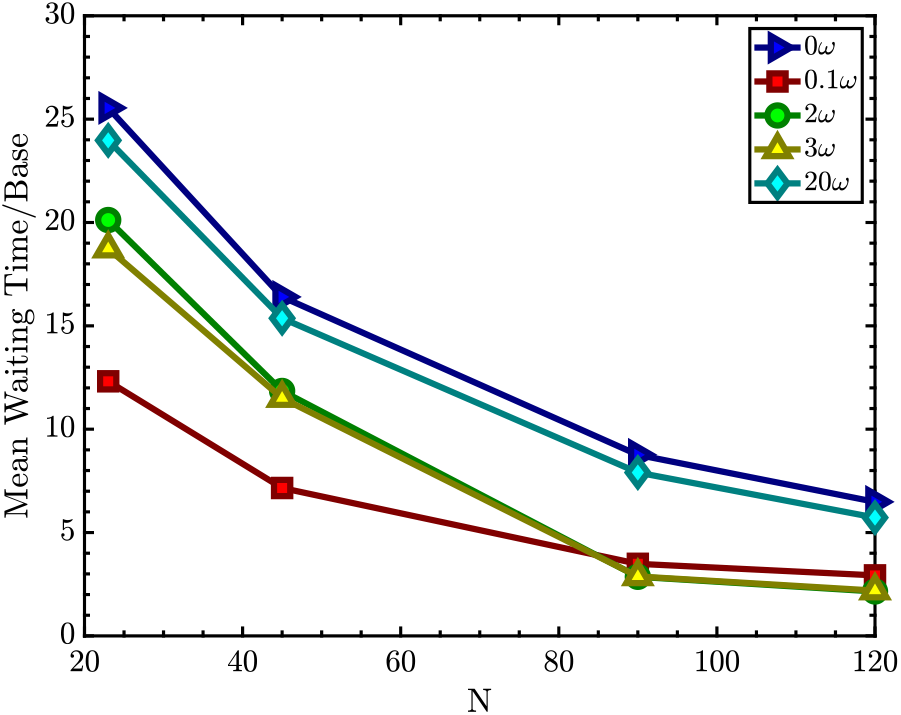
Mean waiting time per base *v.s.* DNA length for different frequencies.

## IV. CONCLUSIONS

We have looked in detail at the translocation dynamics of DNA in the presence of oscillating field. During the positive amplitude of the oscillation the system facilitates DNA translocation, and during the negative amplitude of the oscillation the system hinders translocation. Translocation time is lower at very low oscillating frequencies, 0.1*ω* to 1*ω*, as a result of the prolonged effect of the facilitating state. These fast translocations are characteristic of system under maximum facilitation. At intermediate frequencies, 2*ω* to 7*ω*, the mean translocation time varies non-monotonically for longer DNA N = 90 and 120 but increases steadily for shorter DNA, N = 23 and 45. The different dependence of translocation time with frequency for different chain lengths indicate that stochastic resonance is length dependent. For N = 90 the SR frequency is at 3*ω* where a maximum in the translocation time is observed. For N = 120 the SR frequency is at 2*ω*. Stochastic resonance is not observed for N = 23 and 45. At high frequencies, 10 and 20*ω*, the translocation time is similar to the zero frequency case as the duration of the facilitating/hindering state is too short for DNA to respond to.

We saw that DNA can extend and contract in response to the oscillating field and the number of peaks in DNA extension scales loosely and qualitatively with the oscillation. The extension/contraction is observed even after DNA has translocated out of the nanopore.

We also analyzed the time DNA spent in the vestibule before translocating. The longest waiting time is seen for zero frequency cases for all DNA lengths. Waiting time shows a non-monotonic dependence with frequency, and also with chain length. Mean waiting times decrease as soon as the oscillating field is applied as DNA motion become strongly facilitated. We showed that waiting time decreases with chain length and for very long DNA waiting time will all be the same regardless of the oscillation frequency.

The observed SR frequency can be readily adopted to real systems. Experiments could be used to fine-tune the SR frequency and conditions. Other types of protein nanopore might also be of great interest to see if SR can be achieved under similar conditions and to compare how changes in the nanopore geometry affects SR conditions.

## ACKNOWLEDGMENTS

This work would not have been possible without the support from the National Science Foundation (DMR-1713696), National Institutes of Health (Grant No. R01HG002776-15), and AFOSR (Grant No. FA9550-14-1-0164). We are also thankful for the helpful discussions with Dr. Zachery Dell.

## Notes

### Competing Interest Statement

The authors have declared no competing interest.

