## Supporting Information for "Stochastic Resonance Behavior of DNA Translocation with an Oscillatory Electric Field"

Ining A. Jou

*Department of Polymer Science & Engineering*

Rhys A. Duff

*Department of Chemical Engineering*

Murugappan Muthukumar*

*Department of Polymer Science & Engineering,*

*University of Massachusetts Amherst.*

*

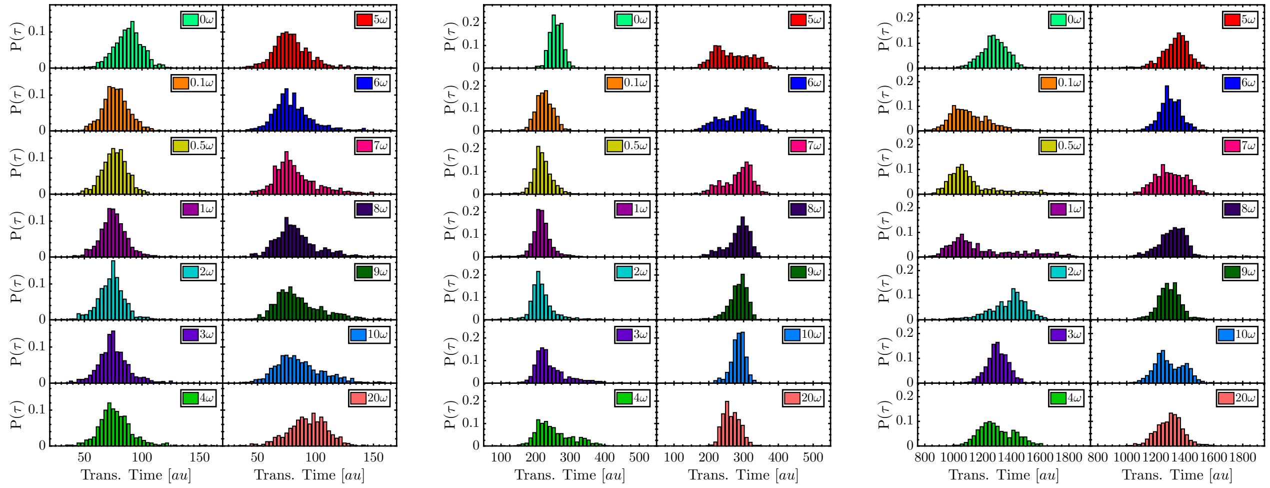


(a) $N = 23$ (b) $N = 45$ (c) $N = 120$

FIG. S1: Probability distribution of DNA translocation time in the presence of various frequencies of the oscillating field.

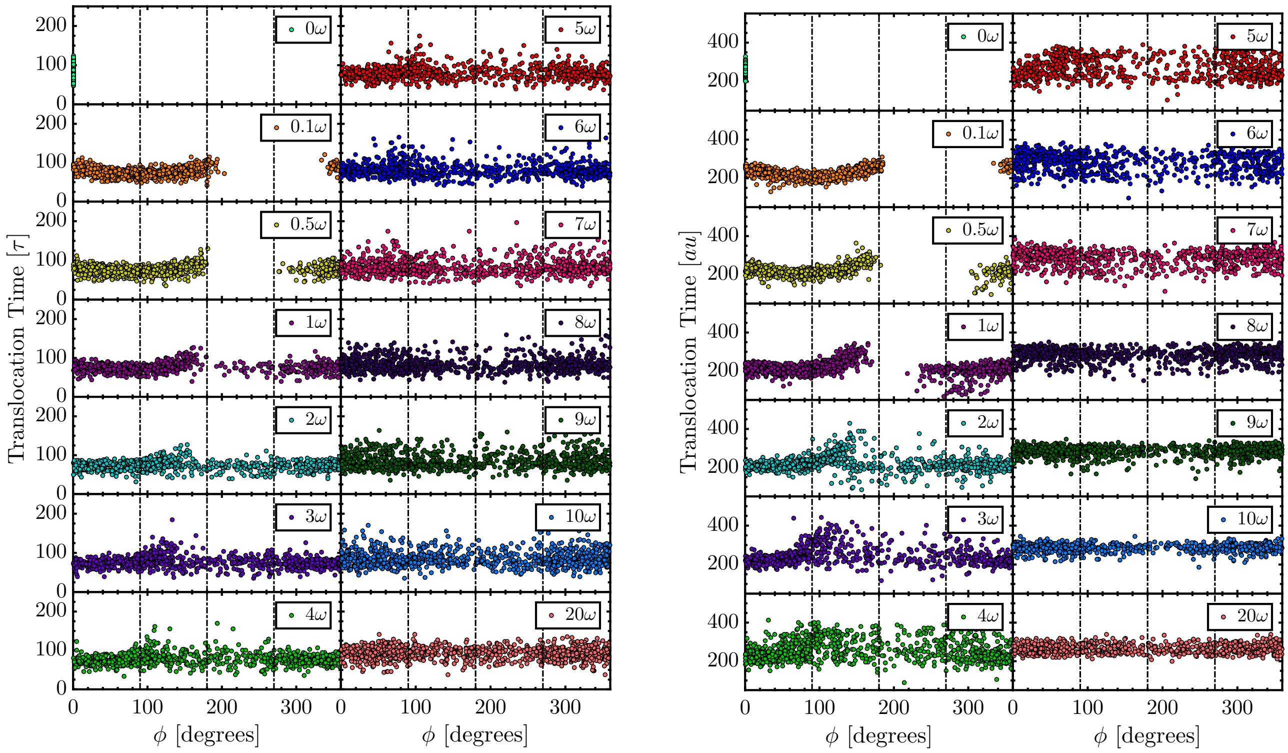


(a) $N = 23$ (b) $N = 45$

FIG. S2: Scatter plot of DNA translocation time *vs*. the randomly generated initial phase $\phi$, where $0^\circ \leq \phi \leq 360^\circ$. Vertical dashed lines are at $90^\circ$, $180^\circ$, and $270^\circ$, respectively.

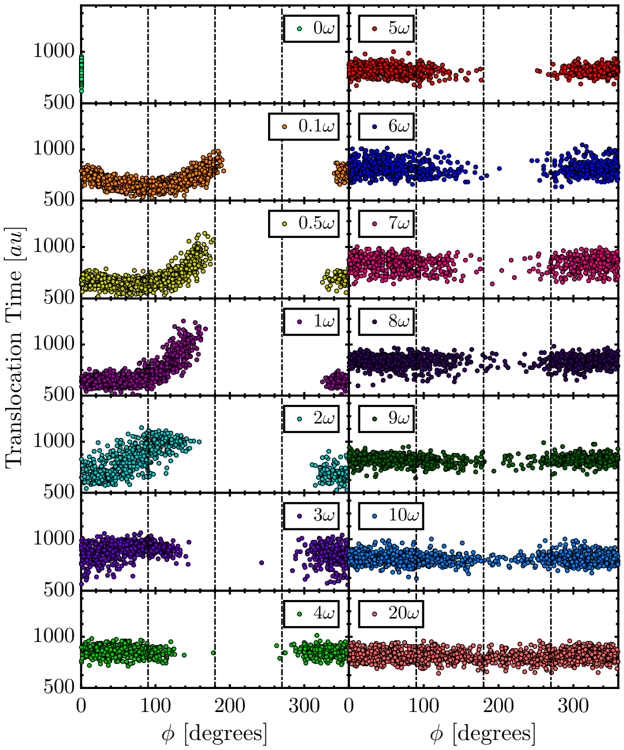


(a) $N = 90$

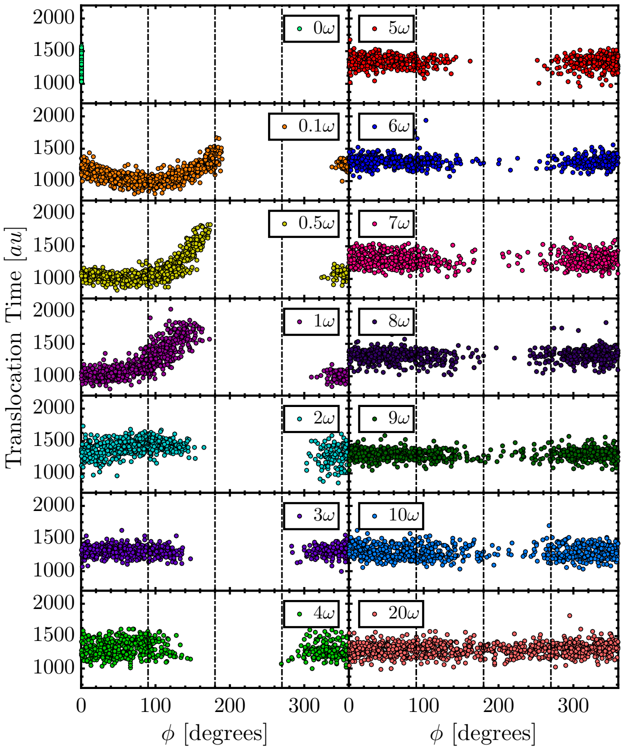


(b) $N = 120$

FIG. S3: Scatter plot of DNA translocation time *vs*. the randomly generated initial phase $\phi$, where $0° \leq \phi \leq 360°$. Vertical dashed lines are at 90°, 180°, and 270°, respectively.

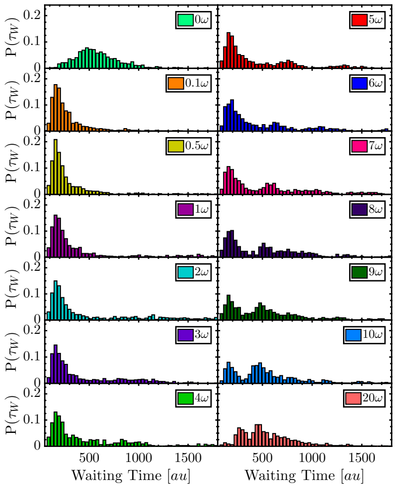


(a) $N = 23$

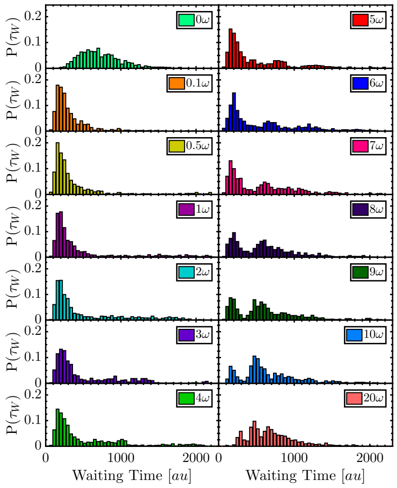


(b) $N = 45$

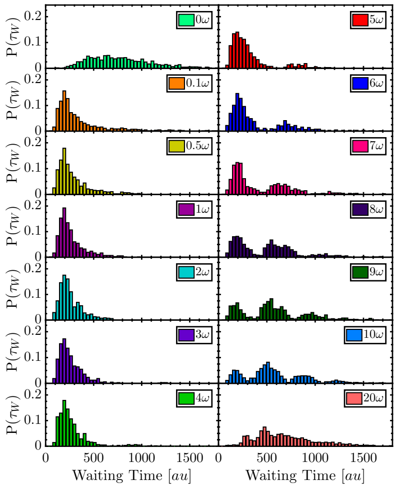


(c) $N = 120$

FIG. S4: Probability distribution of DNA waiting time in the presence of various frequencies of the oscillating field.

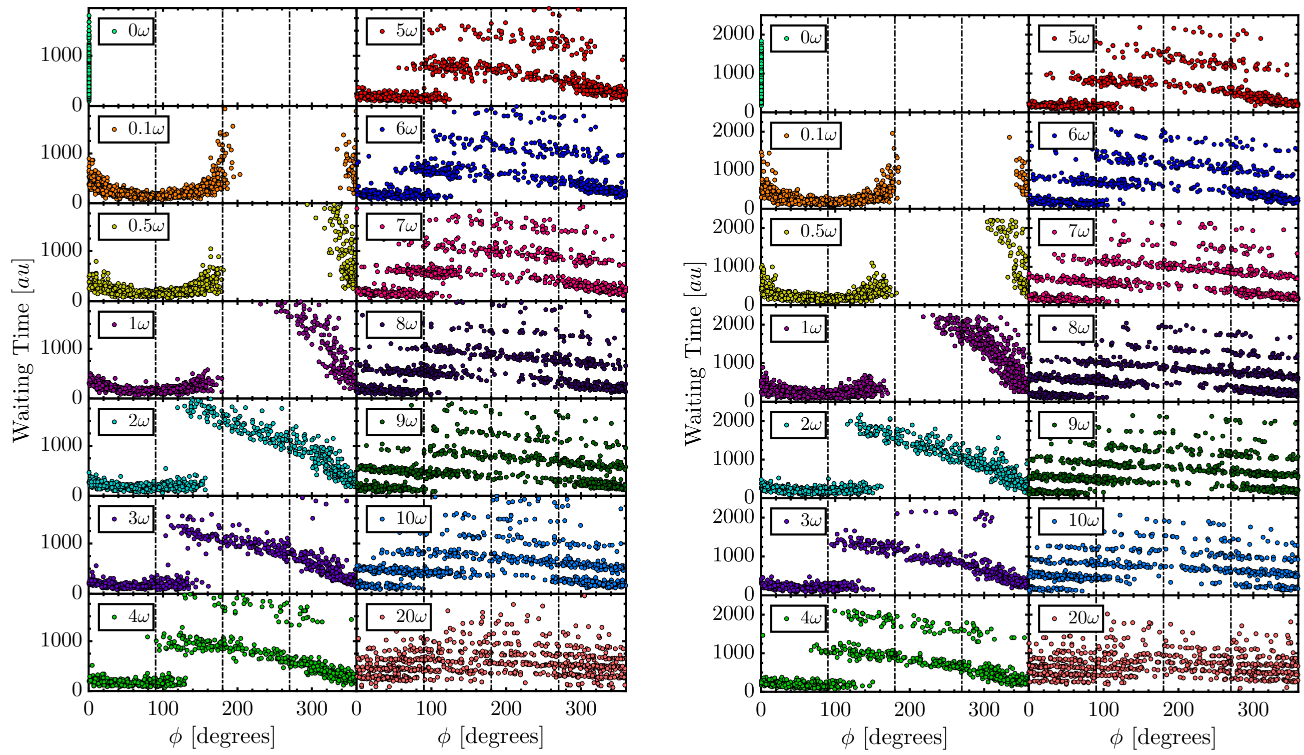


(a) $N = 23$ (b) $N = 45$

FIG. S5: Scatter plot of DNA waiting time *vs*. the randomly generated initial phase $\phi$, where $0^\circ \leq \phi \leq 360^\circ$.

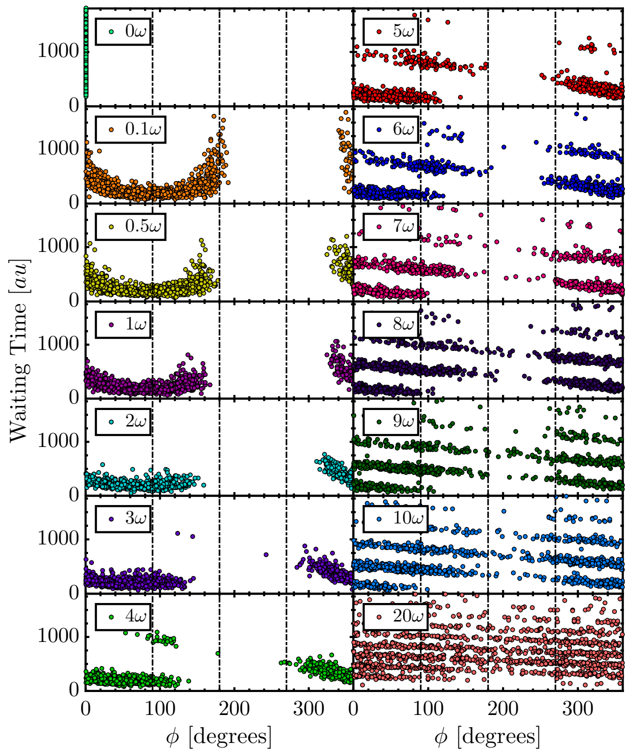


(a) $N = 90$

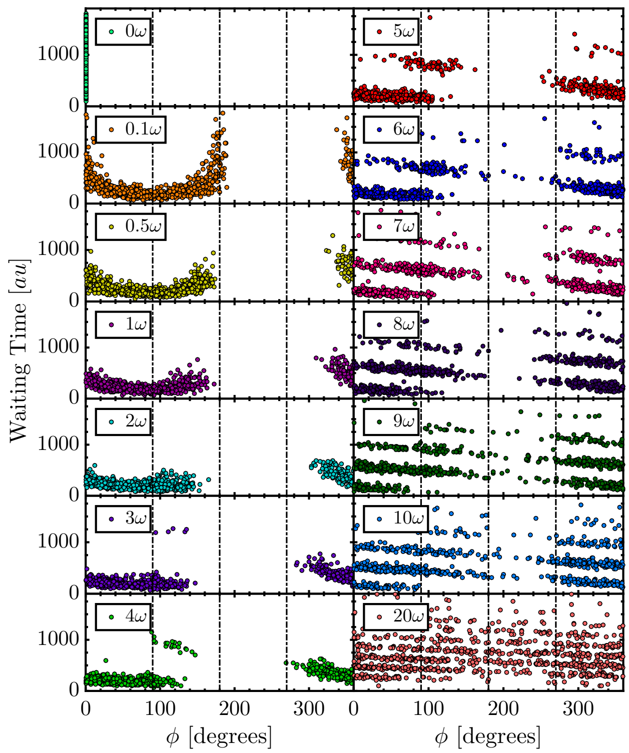


(b) $N = 120$

FIG. S6: Scatter plot of DNA waiting time *vs*. the randomly generated initial phase $\phi$, where $0^\circ \leq \phi \leq 360^\circ$.
